# Efficient neural decoding of self-location with a deep recurrent network

**DOI:** 10.1101/242867

**Authors:** Ardi Tampuu, Tambet Matiisen, H. Freyja Ólafsdóttir, Caswell Barry, Raul Vicente

## Abstract

Place cells in the mammalian hippocampus signal self-location with sparse spatially stable firing fields. Based on observation of place cell activity it is possible to accurately decode an animal’s location. The precision of this decoding sets a lower bound for the amount of information that the hippocampal population conveys about the location of the animal. In this work we use a novel recurrent neural network (RNN) decoder to infer the location of freely moving rats from single unit hippocampal recordings. RNNs are biologically plausible models of neural circuits that learn to incorporate relevant temporal context without the need to make complicated assumptions about the use of prior information to predict the current state. When decoding animal position from spike counts in 1D and 2D-environments, we show that the RNN consistently outperforms a standard Bayesian model with flat priors. In addition, we also conducted a set of sensitivity analysis on the RNN decoder to determine which neurons and sections of firing fields were the most influential. We found that the application of RNNs to neural data allowed flexible integration of temporal context, yielding improved accuracy relative to a commonly used Bayesian approach and opens new avenues for exploration of the neural code.

**Author summary:** Being able to accurately self-localize is critical for most motile organisms. In mammals, place cells in the hippocampus appear to be a central component of the brain network responsible for this ability. In this work we recorded the activity of a population of hippocampal neurons from freely moving rodents and carried out neural decoding to determine the animals’ locations. We found that a machine learning approach using *recurrent neural networks* (RNNs) allowed us to predict the rodents’ true positions more accurately than a standard Bayesian method with flat priors. The RNNs are able to take into account past neural activity without making assumptions about the statistics of neuronal firing. Further, by analyzing the representations learned by the network we were able to determine which neurons, and which aspects of their activity, contributed most strongly to the accurate decoding.

## Introduction

Place cells, pyramidal neurons found in CA1 and CA3 of the mammalian hippocampus [1–4], exhibit spatially constrained receptive fields, referred to as place fields. In general, the activity of place cells is considered to be stable [5, 6]; place fields are typically robust to the removal of specific environmental cues [7,8], persist between visits to a location [9], and in the open field do not strongly depend upon an animal’s behaviour [2, 5]. Upon exposure to a novel enclosure the firing correlates of place cells rapidly ‘remap’; place fields change their firing rate and relative position, forming a distinct representation for the new space [10–12]. For these reasons place cells are widely held to provide the neural basis of self-location, signalling the position of an animal relative to its environment and thus being a necessary element for the control of spatial behaviours, such as navigation, and the retention of spatial memories [2]. Unsurprisingly then, given information about the activity of a population of place cells, it is possible to decode the location of an animal with a relatively high degree of accuracy [13, 14].

However, although place cell activity is strongly modulated by self-location this relationship is non-trivial and not exclusive. For example, during rest and brief pauses, but also during motion, the place code can decouple from an animal’s current location and recapitulate trajectories through the enclosure [15]; ‘replaying’ previous experience [16] or, perhaps, foreshadowing future actions [17]. Similarly, when animals run on linear runways or perform constrained navigational tasks, such as T-maze alternation, place cell activity becomes strongly modulated by behaviour, disambiguating direction of travel [18], prospective and retrospective trajectories [19,20], and the degree of engagement with a task [21]. Furthermore, although place fields are repeatable they are not static. Even though remapping occurs rapidly in a novel environment, the newly formed firing fields continue to be refined during subsequent experience, a process that appears to persist for several hours [10, 13, 22, 23]. Even in familiar environments, that animals have visited many times, the spatial activity of place cells is known to exhibit incremental changes that can result in the generation of distinct spatial codes [23–26], which might be important for encoding goal locations [27] or other non-spatial variables [28]. As such, although hippocampal activity provides considerable information about an animal’s self-location the representation is dynamic: accumulating changes and sometimes encoding other variables both spatial and non-spatial.

A common approach used to interrogate neural representations, such as that of place cells, is decoding; the accuracy with which a variable, such as self-location, can be decoded from the brain, places a useful lower limit on the amount of information present [13, 14]. In the case of place cells, decoding methodologies typically apply a Bayesian framework to calculate a probability distribution over the the animal’s position, given the observed neural data [14, 16]. Decoding to a specific location is then accomplished via a maximum likelihood estimator applied to the probability distribution. However, the accuracy of Bayesian methods depends on accurate information about the expected activity of neurons. For place cells, activity recorded over the course of tens of minutes is typically used to estimate the firing rate of each cell at different points in the animal’s enclosure, with instantaneous rates assumed to exhibit Poisson dynamics. However, for the reasons outlined above, it is not clear that hippocampal activity can be modelled in this way. Indeed, the variability of place cell firing rates is known to greatly exceed that expected from a Poisson process [29]. As such, it is likely that Bayesian methods, as currently applied, do not provide an accurate reflection of the accuracy with which the hippocampus encodes self-location.

To better understand these constraints, we trained a deep recurrent neural network (RNN) [30–32] to decode rodent location from the firing rate of CA1 neurons. At each time step the network was presented with a vector corresponding to the spike counts of hippocampal cells within a given time window. After accumulating information for 100 time-steps the network was required to predict the animal’s location – supervision being provided in the form of the animal’s true location. We found that decoding with the trained RNN was consistently more accurate than a standard Bayesian approach [14,16]. This demonstrates that RNNs are able to capture the relationship between a temporal sequence of neural activity and an encoded variable without the necessity of explicit assumptions about the underlying noise model or complicated hand-coded priors. Further, inspection of the trained network allowed us to identify both the relative importance of individual neurons for accurate decoding and the locations at which they were most informative. Thus, not only does the accuracy of the RNN set a new limit for the amount of information about self-location encoded by place cells but more generally this work suggests that RNNs provide a useful approach for neural decoding and provide a means to explore the neural code.

## Results

### High accuracy decoding of self-location in 2D environments

To test the RNN’s ability to decode rodent location based on hippocampal activity we first characterized the decoding error for a single animal foraging in a 2D arena (1m x 1m square). Single unit recordings were made using tetrodes from region CA1 of five rats. Rat R2192 yielded the greatest number of simultaneously recorded hippocampal neurons (n=63). Since the number of recorded neurons is expected to correlate with decoding accuracy, we first focused on this particular animal.

Neural data was processed to extract action potentials and these were assigned to individual neurons using the amplitude difference between tetrode channels [33] (see Methods). The input features for the RNN-decoder then consisted of spike counts for each neuron within a set of time windows. The length of time windows was parametrically varied between 200 ms and 4000 ms in 200 ms increments. Each consecutive window started 200 ms later than previous one (this means 0% overlap for 200 ms windows, 50% overlap for 400 ms windows, 80% overlap for 1000 ms windows, etc. See ”Feature extraction” in Methods). The network was presented with spike counts from 100 windows before being asked to predict the animal’s location at the center of the latest window.

As the RNN training process is stochastic, 10-fold cross validation (CV) procedure was run multiple times for each window size. For each of these runs we trained 10 models (for each fold of CV) and extracted the mean and median results across the folds. Black dots on Fig 1 correspond to these different realizations of the 10-fold CV procedure (notice multiple dots per window size). 10-fold cross validation was also applied to the Bayesian decoder.

**Fig 1.**
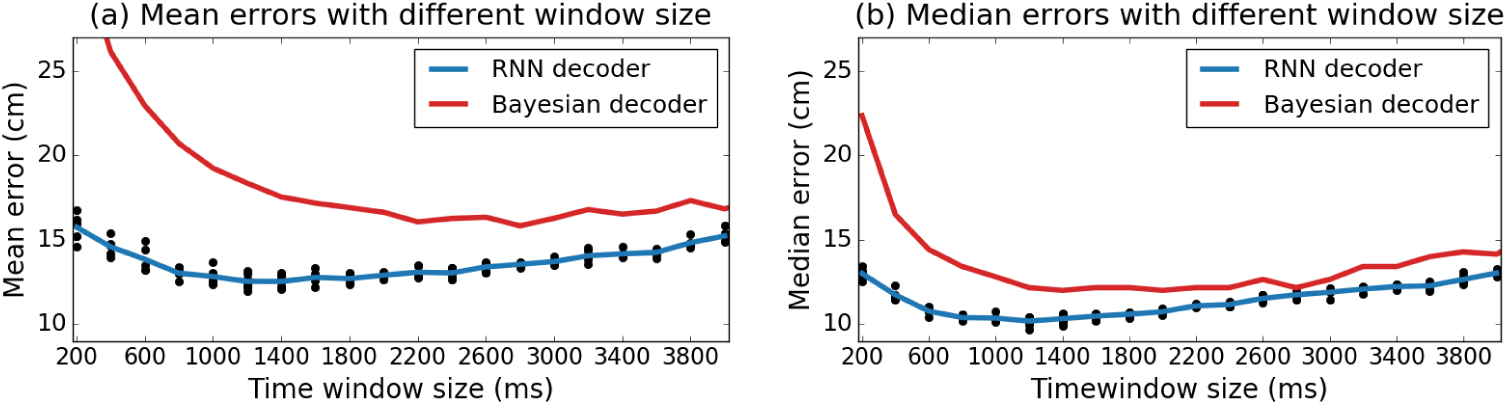
Accurate decoding of position with a RNN. Location decoding errors based on CA1 neural data recorded from 1m square open field environment as a function of time window size (mean error in left panel, median error in right panel). Blue lines represent errors from the RNN decoder and red lines from a Bayesian approach. Results for the RNN approach are averaged over different independent realizations of the training algorithm. Black dots depict the mean/median error of each individual model. Results shown are for animal R2192.

For both the mean (Fig 1a) and median (Fig 1b) of the validation errors, the error curve was convex with lowest errors obtained at intermediate values. Best median decoding accuracy was achieved with time window of 1200 ms (median error = 10.18 ± 0.23 cm). Best mean decoding was achieved for a timewindow of 1400 ms (mean error = 12.50 ± 0.39) cm). Using longer or shorter time windows lead to higher errors – likely because spike counts from shorter windows are increasingly noisy, while the animal’s CA1 activity is less specific to a particular location for longer windows. For all time windows, the accuracy of the RNN considerably exceeded that of the Bayesian decoder (red line). The lowest median decoding error with Bayesian decoder was 12.16 cm (19.5% higher than for the RNN; this best accuracy could be obtained with multiple different window sizes for Bayesian), lowest mean error was 15.83 cm with 2800 ms windows.

The RNN has the ability to flexibly use information from all 100 input vectors and thus integrates contextual information over time. This results in lower mean and median errors as compared to a baseline Bayesian approach that does not have access to information about past activity. In particular, that RNN approach achieves its best results for shorter time windows than the Bayesian approach. We hypothesize that having access to contextual information helps to overcome the stochastic noise in the spike counting obtained for shorter time windows.

Beyond the global descriptors of mean and median error, we also inspected the distribution of decoding error sizes (Fig 2a). For the RNN the distribution followed a unimodal curve with most predictions deviating from the rat’s true position by 6–8 cm. Few errors were larger than 35 cm (1.7 % of errors > 35 cm). The simple Bayesian classifier achieves more very low (< 2 cm) errors, but also an abundance of very large (> 50 cm) errors (7.9 % of errors > 35 cm, 2.8 % > 50 cm).

**Fig 2.**
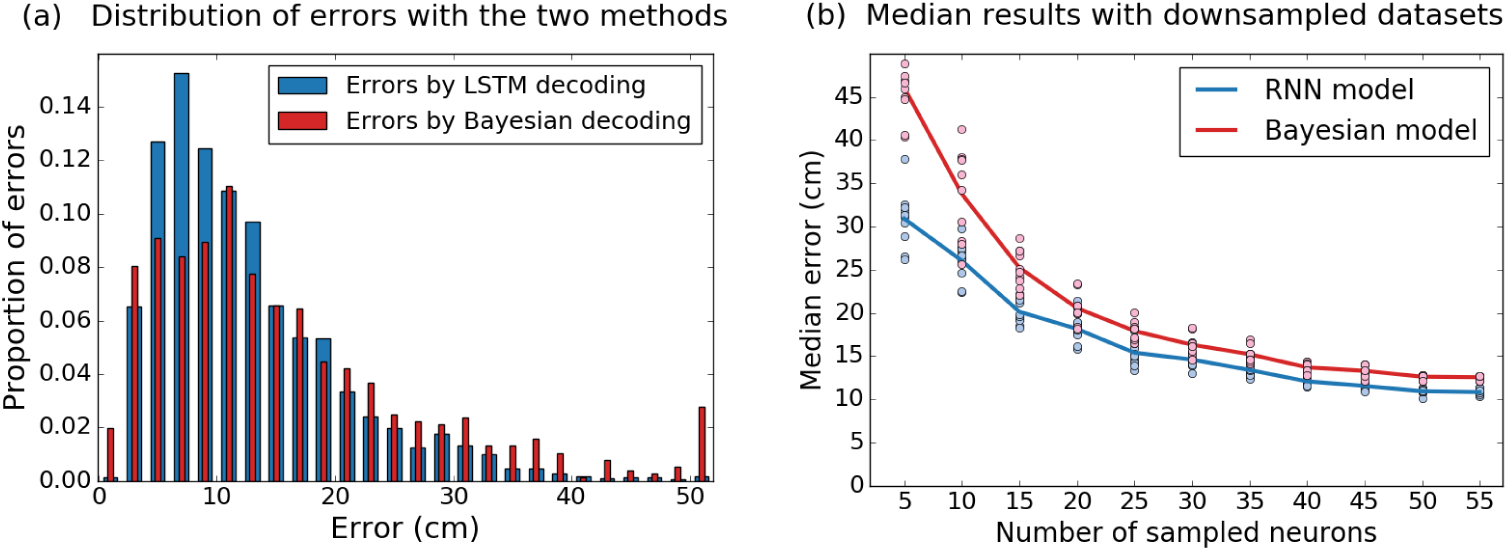
Comparison of RNN and Bayesian decoders. (a) Histogram of error sizes, generated in each case with the best performing time window (1400 ms for RNN, 2800 ms for Bayesian). The Bayesian decoder makes more very large errors (0.02% vs 2.8% of errros > 50 cm). Errors are grouped into 2 cm bins, the last bin shows all errors above 50 cm. (b) Downsampling analysis demonstrates the RNN decoder is more robust to small dataset sized. Data from R2192 was downsampled such that both decoders were trained with a random subset of the available neurons. For each sample size, 10 random sets of neurons were selected and independent models trained as before using 10-fold cross validation. Dots represents median error for each downsampled dataset. Lines indicate the mean over sets of the same size.

In many cases single unit recordings yield fewer than the 63 neurons identified from R2192. We hypothesised that the RNN’s ability to use contextual information would be increasingly important in scenarios where neural data was more scarce. To test this prediction we randomly downsampled the dataset available from R2192, repeating the training and decoding procedure for populations of neurons varying in size from 5 to 55 in increments of 5. As expected we saw that decoding accuracy reduced as the size of the dataset reduced. However the RNN was considerably more robust to small sample sizes, decoding with an error of 30.9 cm with only 5 neurons vs. 46.0 cm error for the Bayesian decoder (Fig 2b).

### Population-level results in 2D and 1D environments

In total we analyzed recordings gathered during 2D open field free foraging task from five animals (1m x 1m square). For each of these 5 datasets, we determined the best performing time window size for the RNN and Bayesian decoder (similarly to Fig 1). The optimal time window sizes for the five 2D foraging datasets are given in Table 1 along with the length of the recording and the number of identified neurons.

**Table 1.**
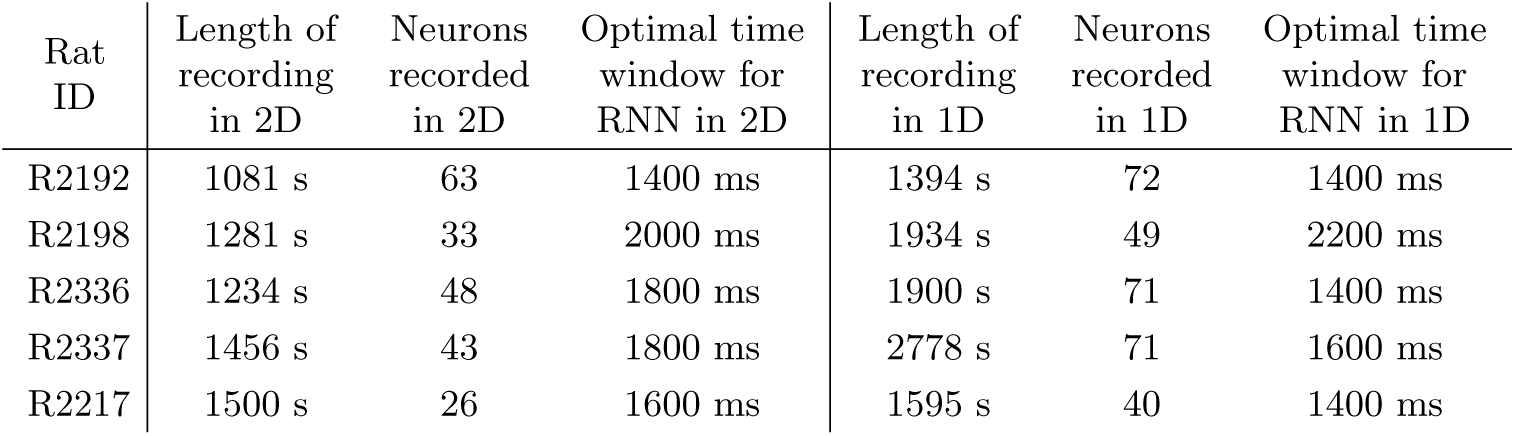
Datasets for 2D and 1D decoding tasks. Number of data points, number of recorded neurons, and the optimal time window for the RNN decoder for each of the 5 analyzed animals and for both decoding tasks.

In the 2D decoding task, for different animals, the mean error across cross validation folds ranges between 12.5-16.3 cm and median between 10.3-13.1 cm (Fig 3a-b). Interestingly, despite some recordings yielding as few as 26 or 33 cells, the decoding accuracy using RNNs is roughly similar. In all cases the mean and median decoding results from the 2-layer LSTM RNN outperformed the standard Bayesian approach.

**Fig 3.**
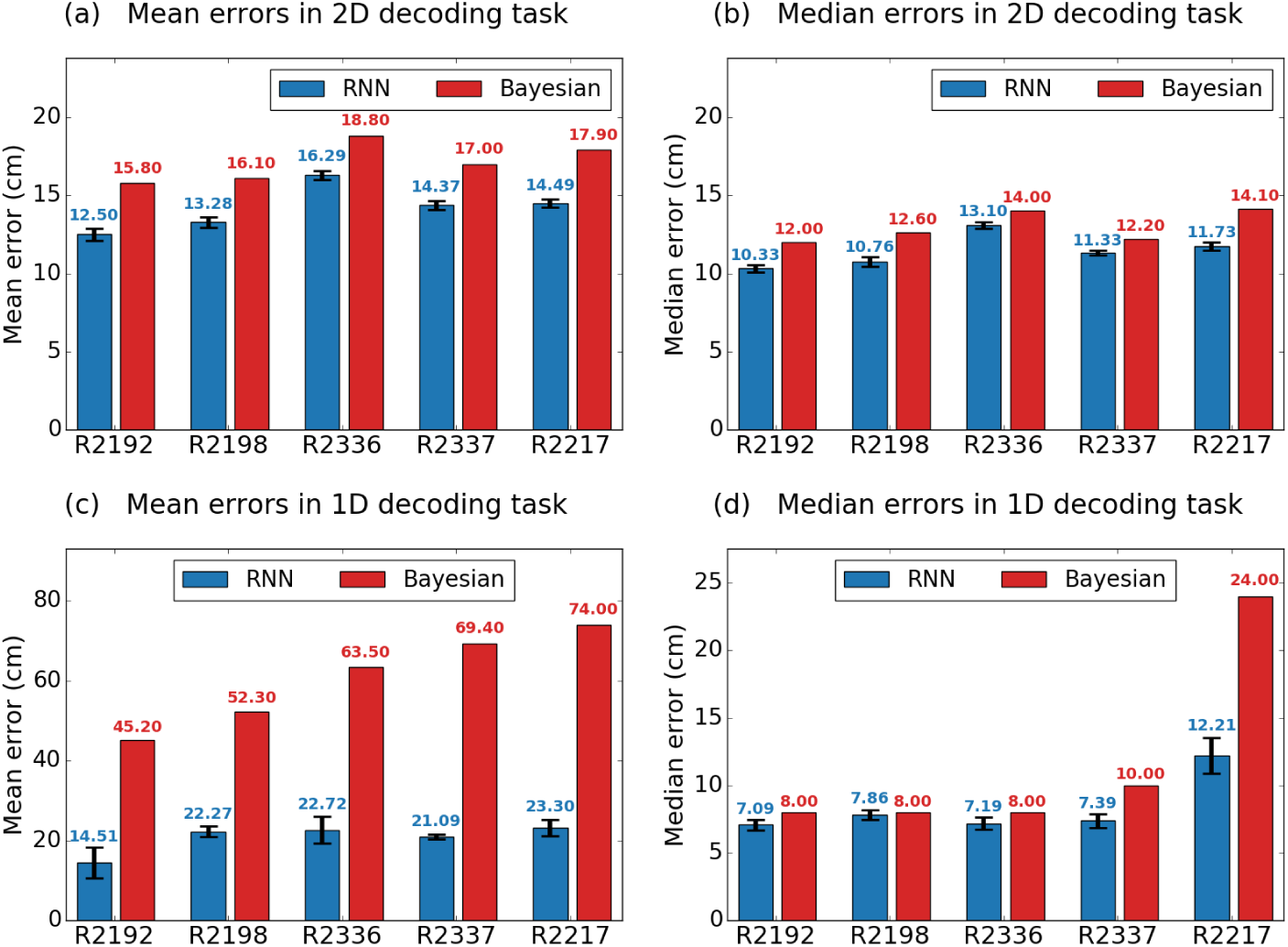
Spatial decoding across animals in 2D and 1D environments. (a-b) Decoding results in a 1m square environment. RNN consistently outperforms the standard Bayesian approach in all 5 data sets. Mean and median errors across cross validation folds, respectively. (c-d) Decoding errors from a 600 cm long Z-shaped track. RNN consistently yields lower decoding errors than a Bayesian approach, the difference is more marked when mean (c) as oppose to median (d) errors are considered.

We also performed decoding on 1D datasets recorded while the same 5 animals shuttled back and forwards on a 600 cm long Z-shaped track for reward placed at the corners and ends (Table 1) [34]. As before we applied RNN and Bayesian decoders to 10-fold cross validated data, selecting in each case the optimal time window size (Table 1). The RNN decoder greatly outperformed the baseline Bayesian decoder in all 5 data sets when comparing mean errors (Fig 3c). However, notice that the Bayesian decoder is a classifier — it is penalized as much for small mistakes as it is for large ones, making it by design more prone to very large mistakes. In the 2D task the largest possible error was 141.7 cm (if the predicted location is in the corner diagonally opposite to the true location), whereas in 1D task it is 600cm (if the opposite end of the track is predicted). In the 1D task a small number of extremely large errors will inflate the mean error, whereas the median will be less affected (Fig 3c-d). Examining the median errors we found that RNN outperformed the Bayesian decoder in all cases. However for four of the five animals the difference in error was relatively small (Fig 3d). For the fifth rat with the fewest cells (R2117, n=40), the RNN clearly outperformed the Bayesian approach, having a median decoding error that was almost half that of the Bayesian decoder.

### Analysis of results obtained with RNN-decoder

Next to understand how behavioural and neural variability influenced decoding accuracy we focused on the results obtained from rat R2192 in the 1m square — the animal with the greatest number of neurons and the lowest decoding error).

First we examined the decoding error as a function of the rat’s location. It is important to note that the animals’ behaviour is non-uniform — the rats visits some parts of the arena more often than others (see Fig 4a). Since more training data is available for frequently visited regions it is expected that any decoding approach would be most accurate in those locations. The spatial distribution of decoding error for R2192 seems to confirm this conjecture — well sampled bins in the center of the enclosure and portions of its borders are more accurately decoded (Fig 4b). To confirm this, we calculated the correlation between the decoding error and the number of training data points located within 10 cm radius of the predicted data point, finding a significant negative correlation (Spearman’s Rank Order, *r* = −0.16, *p_val_* << 0.001, *dof* = 4412).

**Fig 4.**
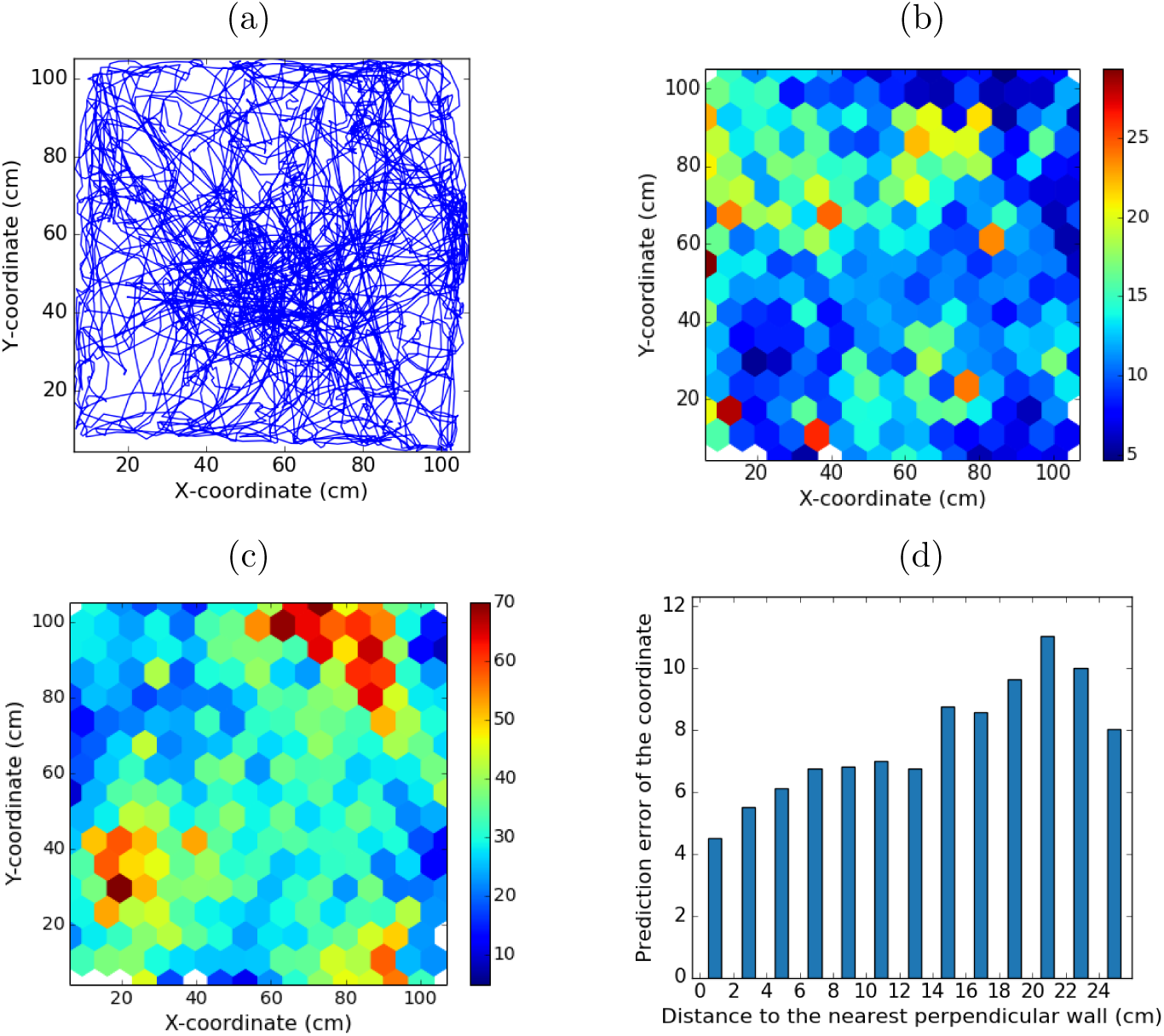
Analysis of the errors in function of location and neural activity for rat R2192. (a) The trajectory of the rat during the entire trial. Not all parts of the arena are visited with equal frequently. (b) The average size of errors made in different regions of space. Color of each hexagon depicts the average euclidean error of data points falling into the hexagon. More frequently visited areas (as seen from (a)) tend to have lower mean error. (c) Sum neural activity in different regions of space. For each data point we sum the spike counts of all 63 neurons in a 1400 ms period centered around the moment the location was recorded. The color of the hexagon corresponds to the average over all data points falling into the hexagon. Areas where sum neural activity is high have lower prediction error. (d) Prediction error of a coordinate decreases if the animal is closer to the wall perpendicular to that coordinate.

Another important factor influencing the decoding accuracy is the distribution of neural activity across the 2D enclosure. In particular, place fields of the recorded hippocampal cells do not cover the enclosure uniformly. Clearly it would be difficult for the algorithm to differentiate between locations where no cell is active. As such, it is likely that areas where more neurons are activated are decoded with higher precision. Our results confirm that the sum of spike counts across neurons at a given location is strongly anti-correlated with the prediction error made at that location (Fig 4c, Spearman’s Rank Order, *r* = −0.31, *p_val_* << 0.001, *dof* = 4412).

We also inspected the *x* and *y* components of the decoding error separately. Previous work suggests that, in the case of grid cells, contact with an environmental boundary results in a reduction of error in the representation of self-location perpendicular to that wall [35]. Such a relationship would be expected if boundaries function as an elongated spatial cue, used by animals to update their representation of self-location relative to it’s surface. Accordingly, we found that for RNN decoding based on CA1 neurons, the decoding accuracy orthogonal to environmental boundaries increased with proximity to that boundary (Fig 4d, Spearman’s Rank Order between error and distance to wall in the region up to 25cm from the wall, *r* = 0.31, *p* << 0.001, *dof* = 3968. The result also held for *x* (*r* = 0.35, *p* << 0.001, *dof* = 2101) and *y* (*r* = 0.25, *p* << 0.001*, dof* = 1855) coordinates separately. Conversely, decoding error parallel to the boundary was not modulated by proximity.

Furthermore, an additional factor that seemed to influence prediction accuracy was the animal’s motion speed. Predictions were more reliable when the rat was moving as opposed to stationary. The mean prediction error for speeds below 0.5 cm/s being 16.5 cm, higher than the 12.1 cm average error for all speeds above 0.5 cm/s (two-sided Welch’s t-test, *t* = 10.62, *p* << 0.001, median errors 8.68 cm and 7.74 cm accordingly). It seems plausible that the lower prediction accuracy during stationary periods might be due to place cells preferentially replaying non-local trajectories during these periods [36]. A second interesting observation is that the prediction error does not increase at higher speeds (two-sided Welch’s t-test between errors in data points where speed is in range from 0.5 cm/s to 10.5cm/s and errors in data points with speed above 10.5cm/s, *t* = 0.31, *p* = 0.76)

### Sensitivity analysis

The accuracy of any neural decoder represents a useful lower bound on the information about the decoded state contained by the recorded neurons. Thus, a biologically relevant question is how such information is distributed among the neurons, across space and time. In short we asked which features of the neuronal activity are the most informative at predicting the animal’s position. To this end we conducted two different types of sensitivity analyses to measure robustness to different types of perturbations.

### Knockout approach

A simple way to estimate the relevance of a specific input in a predictive model is to remove it (to *knock out*) and observe how the prediction accuracy changes. If the input is removed before training, the model can learn to compensate for the missing information — knockout with retraining. However, if the input is removed after training — knockout without retraining — the model cannot adapt or compensate.

Here we used knockout without retraining. The RNN was applied, as before, to predict locations based on a validation dataset in which the activity of a single neuron was set to zero. The knock-out procedure was repeated for each input neuron separately and mean prediction error calculated. Thus we were able to rank neurons by sensitivity - the greater the error increase due to the knocking-out the more crucial the neuron was for the model.

The most influential neuron (neuron #55) was visually identified as an inhibitory neuron based on the lack of clear firing fields and high firing rate 5. In someways it is surprising that this neuron was identified as having the greatest influence on the model — prior work suggests that inhibitory cells do not provide much information about self-location. However, the model’s sensitivity to this neuron is likely due to its high firing rate. Neuron #55 had a firing rate 4 times higher than any other neuron, meaning its removal eliminates the largest number of spikes from the analysis. The other 4 most influential neurons appear to typical pyramidal place cells characterized by clear place fields [2]. The knocking out of these top neurons induced a sizable decrease (> 1*cm*) in the prediction accuracy.

For more than half of the neurons knocking them out decreased the prediction accuracy only very slightly (less than the standard deviation of accuracy, calculated over 10 realizations of the complete model). Among those less influential neurons we found both putative inhibitory interneurons and pyramidal cells with no clear place fields and a lower than average firing rate. For example, the rate map of the least influential neuron, was characterized by a low firing rate #9 (Fig 5f). As suggested by the most and least influential neurons, importance according to knock-out analysis correlated strongly with firing rate (Spearman’s Rank Order, *r* = 0.50, *p_val_* < 0.001, *dof* = 61).

**Fig 5.**
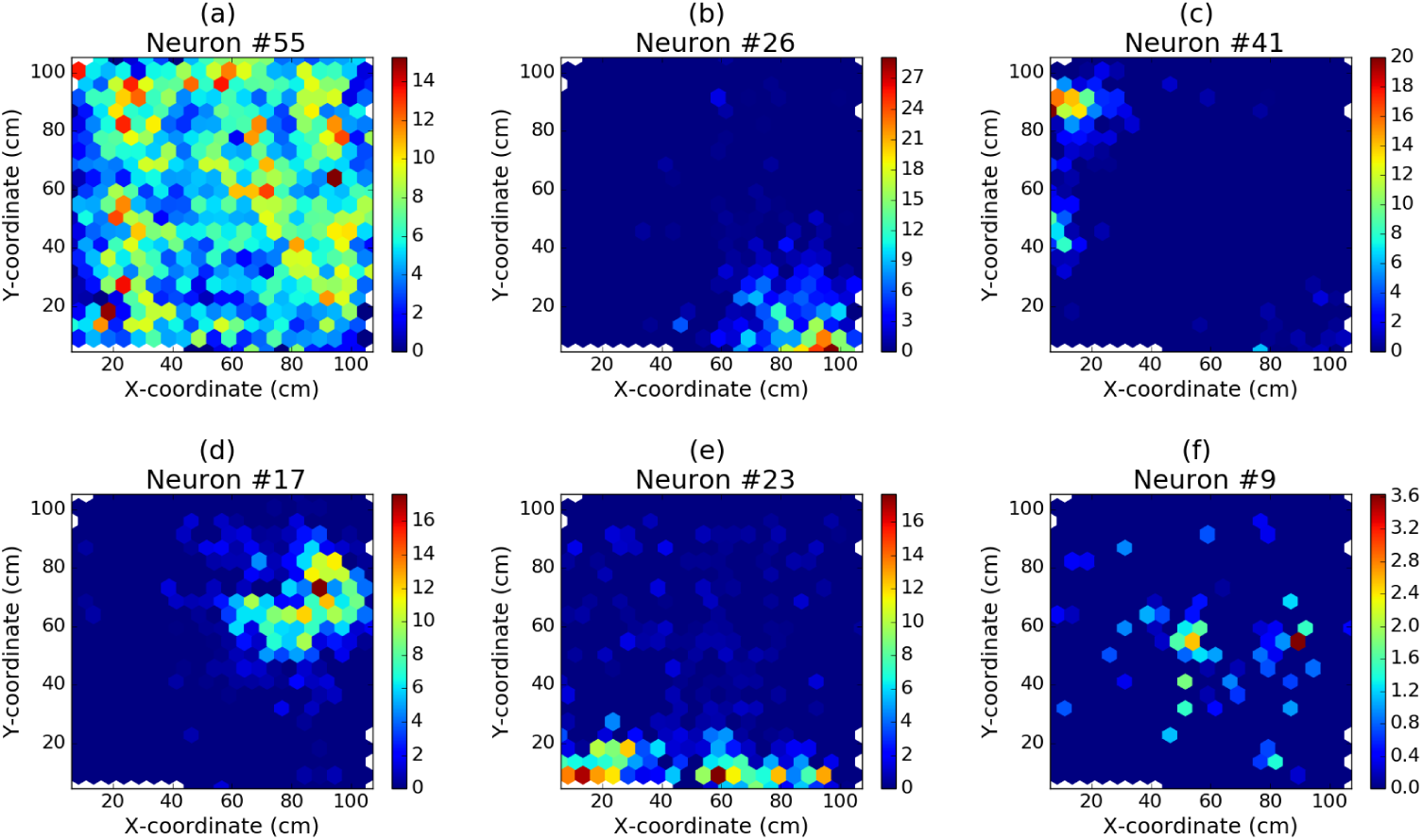
Results of knockout analysis. The firing rate maps of the most (a-e) and least (f) influential neurons according to the knockout analysis. Colour bar to the right of each plot indicates the firing rate in Hz. With the complete dataset the mean error was 12.50 ± 0.28 cm. When knocking out neurons 55, 26, 41, 17, 23 (the five most influential) and 9 (the least influential), the mean error increased to 14.72 cm, 13.80 cm, 13.66 cm, 13.50 cm, 13.49 cm and 12.58 cm respectively.

### Gradients with respect to input

A different way to investigate which neurons most strongly influence decoding accuracy is a gradient analysis. In this analysis we calculate the derivatives of the loss function (mean squared prediction error) of the RNN with respect to the inputs (spike counts of neurons) at different time points. By definition these derivatives show how much a small change in a spike count influences the error. This type of sensitivity analysis is quite different from the knock out analysis — knockout sensitivity measures the impact of silencing a neuron, gradient sensitivity measures the impact of a neurons activity deviating from the expected value.

For each predicted location we asked how sensitive the model was to each of the input spike counts. Since our RNN input is a set of 100 spike count vectors (length of time series), each of length 63 (number of neurons), this amounts to 63×100 gradients per sample. Considering that the whole data set contains around 4400 samples we obtain a 4400×63×100 cubic array of gradient values. To reveal different aspects of the sensitivity of the model, we can average this array of gradients across three dimensions — samples, neurons, or position in the input sequence.

Averaging gradients across all samples and all time windows provided one average gradient value per neuron. Similarly to the knock-out analysis this indicated how relevant the neuron is for the prediction. The two sensitivity measures (knock-out and gradient) were strongly correlated (Spearman’s Rank Order, *ρ* = 0.57, *p* << 0.001, *dof* = 61), but not equivalent. The tests ranked some neurons very differently. For example, the high-firing inhibitory neuron #55 which influences the accuracy most strongly according to knock-out analysis is ranked 47th out of 63 neurons by the gradient sensitivity analysis. Thus illustrating, that despite that fact both measures broadly concur, the gradient and knockout analyses capture different notions of robustness with respect to input perturbations. Interestingly, neither of the two sensitivity measures correlated with the spatial information theoretic measure proposed by Skaggs et al. [37] (Spearman’s Rank Order, gradient-vs-Skaggs: *ρ* = −0.22*, pval* =0.08*, dof* = 61; knock-vs-Skaggs: *ρ* = −0.09, *p* =0.48*, dof* = 61) — likely suggesting that the Skaggs Information Score is not a reliable indication of a neuron’s influence when considered in the context of a population of place cells.

In a second step, we investigate how sensitivity with respect to a neuron’s spike count depends on whether the animal is within its place field or not. Place fields are of variable shape and size and, moreover, a small proportion of the recorded cells have no distinct place fields. Also the firing rates and gradient strengths vary greatly between neurons. Thus, we used firing rate as a proxy to indicate proximity of the animal to a given neuron’s place field — firing rate being maximal when the animal is near the centre of a place field, diminishing the further is moves away from that point. Hence after normalizing both the firing rates and the strength of gradients we averaged over all recorded cells (see the Sensitivity measures subsection in Methods). We saw that sensitivity decreases when the firing rate increases (Fig 6). Hence, indicating that at maximal firing rate — near the center of place field, for example — small changes in firing rate are less influential than they are towards the edges of the firing field. Broadly this accords with theoretical considerations which indicate that, in general, neural responses are most informative in the regions of their coding space where the firing rate changes most rapidly for a given change in the encoded variable [38]. In the case of place cells this corresponds to the edges of the place field. Conversely, outside of the place field, where firing rates fall close to 0 Hz, the sensitivity of the RNN to the neuron is again slightly lower (Fig 6).

**Fig 6.**
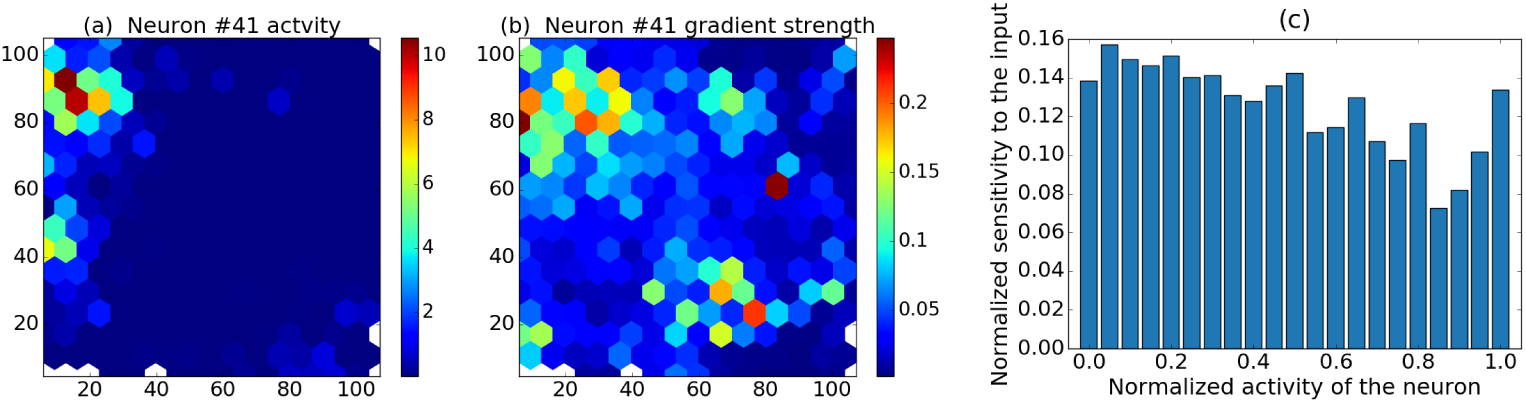
Gradient analysis: sensitivity decreases with activity. (a) Place field of an example neuron. (b) *Sensitivity field* - absolute values of gradients in different locations for the same neuron. (c) Normalized sensitivity as a function of normalized activity across neurons.

## Discussion

We have shown that the sequential processing afforded by an artificial recurrent neural network (RNN) provides a flexible methodology able to efficiently decode information from a population of neurons. Moreover, since a RNN decoder is a neural network, it represents a biologically relevant model of how neural information is processed. Specifically, when applied to hippocampal neural data from freely moving rats [2], the network made use of the past neural activity to improve the decoding accuracy of the animals’ positions. In a 2D open field arena (1m x 1m), the RNN decoder was able to infer position with a median error of between 10.3 cm to 13.1 cm for 5 different rats. These results represented a marked improvement over a standard a Bayesian decoder [14,16] which bases its decision solely on spike counts from a single time window centered around the moment of position measurement. Bayesian methods are known to be optimal decoders when using appropriate priors [39]. However, when applied to neural decoding it is difficult to determine these appropriate priors - as a result sub-optimal approximations are commonly used. Hence we propose that RNNs offer a practical methodology to incorporate sequential context without the need to choose or estimate specific priors over high-dimensional spaces. The improvement in 2D position decoding observed for the RNN was mirrored by similar results from a 1D decoding task using hippocampal recordings made while animals ran on a 6 meter track. Here again, the RNN decoder achieved equal or better results than a standard Bayesian approach.

Making use of the past neural activity as contextual information, the RNN seems more robust to noise than Bayesian classifier. In particular when using shorter time windows the spike counts become noisier and the Bayesian model’s prediction accuracy degraded rapidly. In contrast the RNN decoder was more resistant to the variability of spike counts, likely due to its ability to combine information over the complete sequence of past inputs. Similarly, in situations were fewer neurons were available and hence the total amount information was reduced, the RNN exhibited a pronounced advantage over the Bayesian decoder. Equally, in the 1D task the benefit of the RNN was most evident for animal R2217, which had the fewest recorded neurons. Nevertheless notice that fewer recorded neurons does not necessarily mean lower accuracy. As described in Section 2.3.1, the error depends strongly on the amount of training data available (length of recording) and the quality of the cells (amount and location of firing). Taken together these results suggest that RNN decoding of neural data may prove to be particular useful in situations where large populations of neurons are not available or are difficult to stably maintain.

Beyond quality and amount of data available, the size of error the RNN decoder maed was also seen to depend on the distance of the animal from the walls and its instantaneous speed. At higher speeds (above 10.5 cm/s) the decoding accuracy does not decrease, but when the animal is immobile (below 0.5 cm/s) the error was significantly higher than when in motion. We hypothesize that while stationary hippocampal activity may reflect non-local activity associated with sharp-wave ripple states [36].

Beyond providing more accurate decoding, the neural network approach also provides a new approach to sensitivity analyses. While knockout-type sensitivity analyses can be applied to both Bayesian and RNN decoders, the latter approach also supports gradient analyses. The two types of sensitivity - knockout and gradient - are correlated, but not identical. By design knockout analyses answers how the system behaves if an input is completely removed, while gradient analyses investigated how the system behaves in response to small perturbations to that input. Having access to the gradients with respect to each spike count makes is possible to pose new questions about the dynamic variability of the information content of individual neurons.

## Materials and methods

### Data collection

#### Animals and surgery

Eight male Lister Hooded rats were used in this study. All procedures were approved by the UK Home Office, subject to the restrictions and provisions contained in the Animals Scientific Procedures Act of 1986. All rats (330 − 400*g* / 13 − 18 weeks old at implantation) received two microdrives, each carrying eight tetrodes of twisted 17*µm* HM-L coated platinum iridium wire (90% and 10%, respectively; California Fine Wire), targeted to the right CA1 (ML: 2.2*mm*, AP: 3.8*mm* posterior to Bregma) and left medial entorhinal cortex (MEC) (ML = 4.5*mm*, AP = 0.3 − 0.7 anterior to the transverse sinus, angled between 8 − 10°). Wires were platinum plated to reduce impedance to 200 − 300*k*Ω at 1*kHz*. After rats had recovered from surgery they were maintained at 90% of free-feeding weight with *ad libitum* access to water, and were housed individually on a 12 − *hr* light/dark cycle. MEC data was not analysed for this study.

#### Recording

Screening was performed post-surgically after a 1-week recovery period. An Axona recording system (Axona Ltd., St Albans, UK) was used to acquire the single-units and positional data (for details of the recording system and basic recording protocol see Barry et al(2007). The position and head direction of the animals was inferred using an overhead video camera to record the location of two light-emitting diodes (LEDs) mounted on the animals’ head-stages (50*Hz*). Tetrodes were gradually advanced in 62.5*µm* steps across days until place cells (CA1) and grid cells (MEC) were identified.

#### Experimental apparatus and protocol

The experiment was run during the animals’ light period. First, animals ran on a Z-shaped track, elevated 75*cm* off the ground with 10*cm* wide runways. The two parallel tracks of the Z (190 cm each) were connected by a diagonal section (220*cm*). The entire track was surrounded by plain black curtains with no distal cues. During each track session, animals were required to complete laps on the elevated Z-track. Specifically, the animals were required to run from the start of Arm1 to the end of Arm3, stopping at the track corners and ends in order to receive a food reward. If the animals made a wrong turn at the corners, reward was withheld. Four animals (R2142, R2192, R2198, and R2217) were trained to run on the track for 3 days before recording commenced. For the other animals (R2242, R2335, R2336, R2337), recordings were made from the first day of exposure to the Z-track task. These recordings constitute the dataset we refer to as the 1D decoding task. Not all animals’ recordings were used.

Following the track session the same animals completed a 20min random foraging session in a square (1m x 1m) enclosure. Coverage of the enclosure was encouraged by rewarding animals with sweetened rice. These recordings constitute the dataset we refer to as the 2D decoding task. Not all animals’ recordings were used.

### Decoder based on recurrent neural networks

Deep learning is a class of algorithms that learn a hierarchy of representations or transformations of the data that make the problem of classification or regression easier [30, 32]. In particular, deep neural networks, inspired by biological neural circuits, consist of layers of computational units called neurons or nodes. The deepness means that there are multiple ”hidden” layers between the input and output. By tuning the connection weights between its layers a neural network can learn to approximate a function from a set of examples, i.e., pairs of related input and output data. In this work we are interested in training a neural network to decode the rat spatial coordinates from the activity recorded from its hippocampal cells.

Whereas feed-forward neural networks learn to predict an output based on a single input, recurrent neural networks (RNNs) can deal with series of inputs and/or outputs [31, 32]. In particular, a recurrent network can preserve information from previous inputs by means of feedback connections (loops between its units). Having access to past information can be useful to minimize errors in certain tasks. Such memory of past inputs also means that the order in which the inputs are presented to the network may change the eventual predictions, and thus integrate contextual information over time. A naive implementation of RNNs can only maintain information from a few past inputs, making it possible for the network to detect only immediate trends, but not long timescale dependencies. Advanced realizations of recurrent networks, such as long-short term memory (LSTM) [40] and gated recurrent units (GRU) [41, 42] have specific architecture and sets of parameters that control to what extent past activity should be remembered or overwritten by a new input [42]. This makes them capable of integrating knowledge over a longer sequence. Through using past inputs as contextual information these networks have achieved outstanding performance with noisy sequential data such as text and speech.

#### Network architecture

A RNN can be made to predict (i) a series of outputs based on a series of inputs, (ii) a series of outputs given only one input, and (iii) one output given a series of inputs. For our location prediction task we are interested in the latter - given hippocampal activity (spike counts) over a longer period of time, we aim to predict one set of spatial coordinates - the animal location.

The architecture, illustrated in Fig 7c, of the RNN used in this work consists of an input layer (same size as the number of recorded neurons) followed by two 512-node LSTM layers, and an output layer (2 nodes, one for each spatial coordinate *x* and *y*).

**Fig 7.**
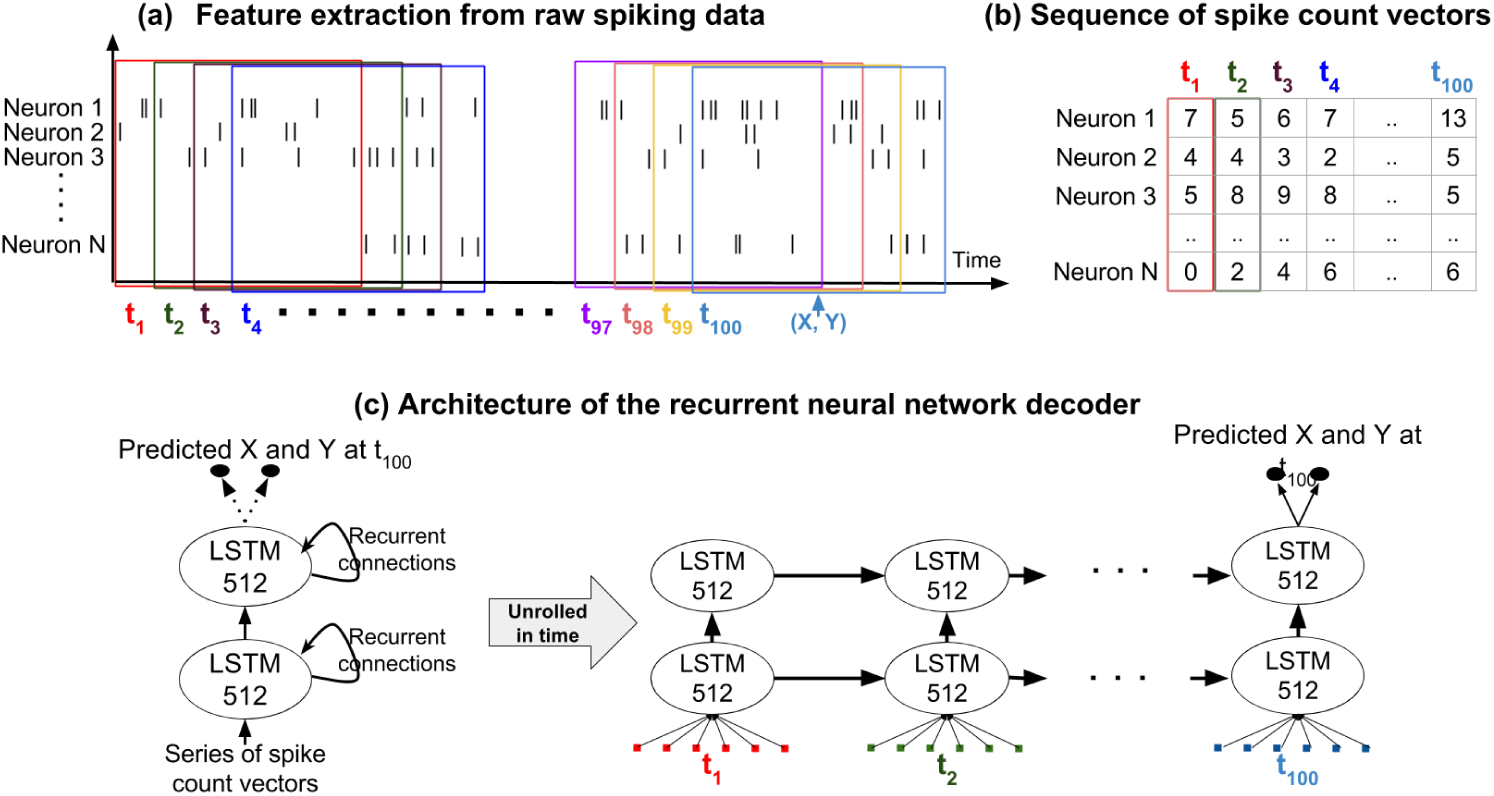
Feature extraction from spiking data and neural network architecture. (a) Extracting a sequence of spike count vectors from spiking data. Each subsequent input originates from an overlapping time period, shifted 200 ms forward in time. (b) The input data that the RNN decoder will use is a sequence of spike count vectors from these time windows. (c) The network used for decoding consists of an input layer (size equals number of recorded neurons), two hidden layers containing 512 long-short term memory (LSTM) units and an output layer of size 2 (*x* and *y* coordinates). The spike count vectors are inserted to the input layer one by one at each timestep. The network produces a prediction for *x* and *y* coordinates only at the end of the sequence, at *t* = 100.

### Feature extraction

The features of neural data used for decoding are the spike counts of all *N* cells recorded (forming a *spike count vector*, as shown in Fig 7a and 7b). In particular, the recurrent neural network is presented with a series of 100 of such spike count vectors, corresponding to activity of all cells in 100 overlapping time windows. The shift between consecutive time windows was fixed to 200 ms for all window sizes (this means 0% overlap in 200 ms windows, 50% overlap in 400 ms windows, 80% overlap in 1000 ms windows). Across the 100 time steps we consider activity from approximately 20 seconds.

Based on this series of 100 spike count vectors the recurrent network was trained to predict the rat’s location in the center of the last (100th) time window. Thus, each sequence of 100 vectors plus the correct location of the rat at the center of the last time window forms one data point for training the RNN.

During the training procedure the network aims to minimize an objective function, in our case the mean squared error of the coordinates. The learning is done for 50 epochs (full cycles of training data) using RMSprop optimizer (variant of stochastic gradient descent), with a mini-batch size equal to 64. All computations were performed with custom-made scripts using Keras neural network library [43].

### Bayesian decoder

Spatial decoding was also implemented using a Bayesian framework [44] subject to 10-fold cross validation (see also the next subsection). Specifically, for each fold, 90% of the data was used to generate ratemaps for hippocampal neurons -spike and dwell time data were binned into 2 cm square bins, smoothed with a Gaussian kernel (*σ*=1.5 bins), and rates calculated by dividing spike numbers by dwell time. Note, for the Z-maze only, positional data was linearised before binning.

Next, with the remaining 10% of the data, using temporal windows (200 ms to 1000 ms) each of which overlapped with its neighbours by half, we calculate the probability of the animal’s presence in each spatial bin given the observed spikes – the posterior probability matrix [14, 16].

Specifically during a time window (T) the spikes generated by *N* place cells was *K* = (*k*_1_,…,*k_i_*,…,*k_N_*), where *k_i_* was the number of spikes fired by the *i* − *th* cell. The probability of observing *K* in time T given position (*x*) was taken as:

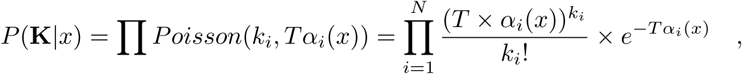
where *x* indexes the 2 cm spatial bins defined on the Z-track/foraging environment and *α_i_*(*x*) is the firing rate of the *i* − *th* place cell at position *x*, derived from the ratemaps.

To compute the probability of the animal’s position given the observed spikes we applied Bayes’ rule, assuming a flat prior for position (*P* (*x*)), to give:

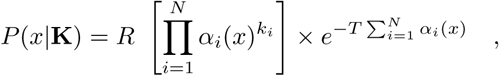
where *R* is a normalizing constant depending on *T* and the number of spikes emitted. Note we do not use the historic position of the animals’ to constrain *P*(*x*|*K*) thus the probability estimate in each *T* is independent of its neighbours. Finally, position was decoded from the posterior probability matrix using a maximum likelihood method - selecting the bin with the highest probability value. Decoding error was then taken as the Euclidean distance between the centre of the decoded bin and the centre of the bin closest to the animal’s true location.

### Cross validation and averaging of results

The reported errors for both Bayesian and RNN approach are measured using a 10-fold cross validation method that divides the *D* data points between training and validation sets. Due to the overlap between consecutive time windows a random assignment of data points to training and validation sets would imply that for most of the validation data points a highly correlated neighbouring sample can be found in the training set. This would result in an artificially high validation accuracy that does not actually reflect the model’s ability to generalize to new, unseen data.

Instead, in our analysis the first fold in cross validation simply corresponds to leaving out the first 10% of the recording time and training the model on the last 90% of data. The second fold, accordingly, assigns the second tenth of recordings to the validation set, and so on. For RNNs we need to additionally discard 99 samples at each border between training and validation sets. Remind that the input for RNNs is a series of 100 spike count vectors - to avoid any overlap between training and test data we remove validation data points that have at least one shared spike count vector with any training data point.

For each fold we train a model on the training set and calculate the error on the validation set. All reported errors are the validation errors - errors that the models make on the one tenth of data that was left out of the training procedure. To increase the reliability of the results, we perform 10-fold cross validation procedure multiple times and report the mean and median of the errors. This is done only for the RNN decoder, because the Bayesian decoder is deterministic and repeating cross-validation procedure multiple times is not necessary.

### Analysis of decoding results

#### Quantifying prediction errors

When decoding rat locations in the 2D arena, prediction errors in the animal position were quantified by the mean Euclidean distance (MED) between the predicted and true positions:

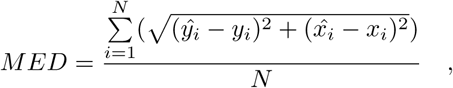
where 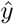 and 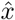 are the locations predicted by the decoder, *y* and *x* are the true locations and *N* is the number of data points.

The training procedure of recurrent neural networks is stochastic and always ends up with slightly different solutions. We repeat the 10-fold cross-validation 10 times, giving us 10 independent predictions for each data point. We report the average of errors over these 10 realizations (and not the error of the averaged prediction).

For evaluating the *x*-coordinate (*y*-coordinate) errors only the *x* (*y*) component of the positions were used in the above formula:

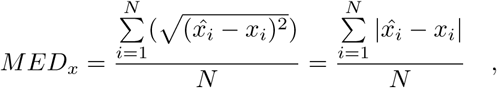

In an additional experiment, we also decode the rat locations on a 600 cm long Z-shaped track. The position of the rat along the track is considered as a 1D coordinate ranging from 0 in one end of the track to 600 in the other end of the Z-shape. To obtain these 1D coordinates the actual locations extracted from camera images are projected to the nearest point on a Z-shaped ideal trajectory. The prediction error of the model is quantified by absolute distance between the predicted and true position along this 1D coordinate.

#### Sensitivity measures

In knock-out analysis we set the activity of a neuron to zero in all validation data points and then calculate the validation errors. The activity is not annulled during training of the model, so the system can not learn to compensate or adapt. We repeat this knock-out procedure for each neuron one by one. We compare how much the prediction error increased when different neurons were knocked out.

The gradient of the loss function with respect to inputs was calculated using back-propagation through time [45], similarly to how gradients with respect to weights are found. Indeed, for updating the connection weights of the network at training time, the algorithm needs to calculate the gradients of the loss function with respect to the weights [30]. These gradients tell us how a small change in a particular weight would influence the final output error. In here we ask a similar question - how much would a small deviation in a certain input change the final loss. Important is to notice that when talking about sensitivity we disregard the sign of the gradient, in all results we use the absolute values (magnitudes) of gradients. We compute gradient strengths for each validation set data point and separately for each neuron’s spike count for each position in the time series of *T* inputs (*T* = 100). This results in a *D* × *N* × *T* matrix of gradient values. To draw further conclusions from the gradient values, we need to average or manipulate this 3D matrix along different dimensions. For example, when calculating the neurons that the model is most sensitive to, we need to average across all data points and all time steps, so we are left with one value per neuron.

When investigating the relationship between sensitivity and location on the place field (on Fig 6c), we also need to normalize the spike counts and gradient magnitudes of different neurons, so that we could aggregate them. To do this one would usually divide the spike count with the maximum value, resulting in measures between 0 and 1 for all cells. In the case or low-firing neurons, however, the noisiness of the data means that the maximum value can be an outlier (we can have maximum count of 4, whereas no other value is above 2). We therefore choose to divide the spike counts with the 99th percentile of the spike count values instead. A few values end up being above 1, but the normalized value distributions of low and high firing neurons look more similar. We do a similar 99-th percentile normalization on the absolute values of gradients. For each normalized firing rate we have one corresponding normalized gradient size. We can then plot how the normalized gradient size depends on normalized firing rate.

## Supporting information

**S1 Text. Temporal gradient analysis**. A third way to investigate the gradients is to average only across the samples. We thus obtain an averaged gradient for each neuron at each different position in the input sequence of 100 time windows. These averages reveal, for example, how sensitive the model is to changes in spike counts of the same neuron at different points of time. Unfortunately recurrent network architecture and training procedures favour information contained in more recent inputs (due to vanishing gradients further back in time). We therefore judge that it is not fair to draw conclusions from comparing sensitivity to spike counts at different positions in the sequence - inputs in the later time steps would show up as more important not necessarily due to information content but due to the algorithm we used. It is however fair to compare the contributions of different neurons at the same time step. We propose to compare the model’s sensitivity to a certain spike count with average of sensitivity across all neurons at the same point of the temporal context sequence. Intuitively such gradient analysis reveals if neuron N’s activity at time window T within the temporal context, was more informative than the activity of other neurons at that time point. This comparison is not distorted by the network architecture, because inputs from different neurons are treated symmetrically (order of neurons could be changed) by the network. No bias exists with respect to either particular neurons or data samples.

As a summary of the analysis described above, Fig 2 shows the normalized gradients of several neurons at different positions within the temporal context. The analysis reveals different profiles of relative sensitivity within the temporal context. In particular, we note that several neurons have a peak in their normalized sensitivity around one second before the last time window for which the animal position is predicted. Nevertheless, our time windows last 1400ms and therefore the temporal resolution is very low. We restrain ourselves from drawing conclusions from this analysis due to lack of temporal precision. We believe that when using smaller, non-overlapping time windows, this type of investigation can reveal interesting temporal aspects of information processing in the brain.

**Fig 1. S1 Fig.**
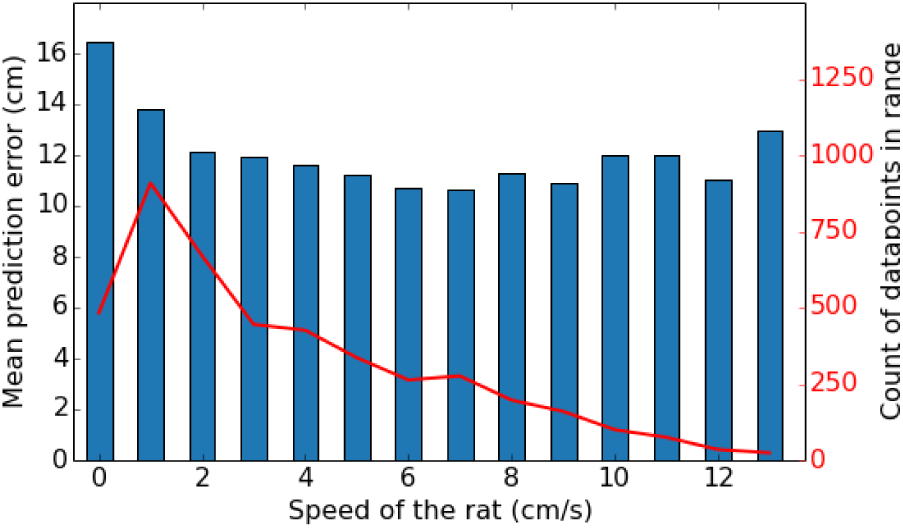
Mean prediction error for different instantaneous movement speeds. Movement speed is based on the distance covered in 200 ms. The first bar is the average over the errors for speeds in range [0, 0.5] cm/s, the second for (0.5, 1.5] cm/s, etc. The error is highest when the rat is not moving or moving very slowly. Notice that speeds in the range of 1-2 cm/s can also be the results of head movements. At higher speeds the exact velocity does not seem to influence accuracy. Note that the bars do not contain the same amount of data points. Apparent changes in the mean error at higher velocities can be attributed to noise as we have less data points there.

**Fig 2.**
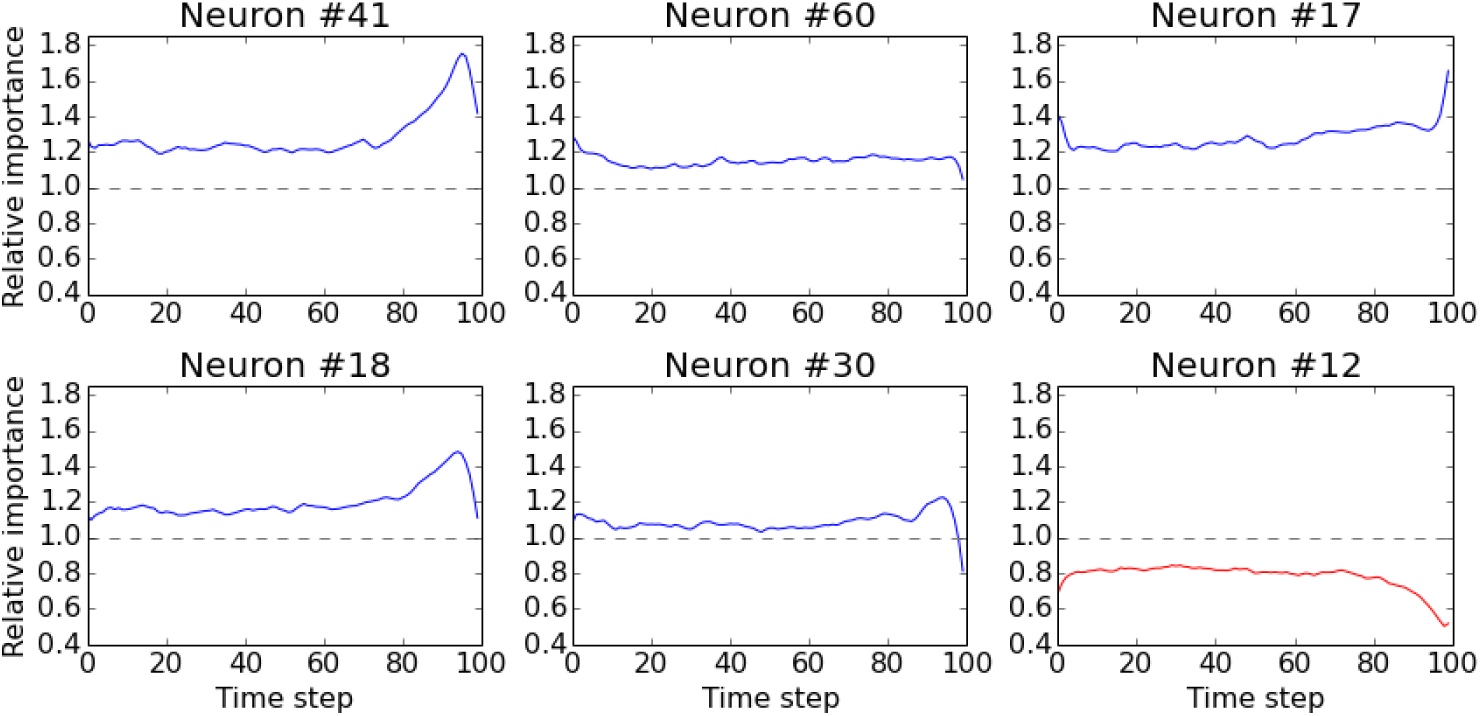
Gradient analysis. a-e) Temporal profiles of relative importance for 5 selected neurons among the highest contributing neurons according to gradient analysis. Notice that the profiles peak at different time steps. f) Temporal profile of the least important neuron according to gradient analysis.

## Acknowledgements

We thank Zurab Bzhalava, Sander Tanni and Jaan Aru for early work and useful discussions. We also thank Jack Kelly for useful discussions. R.V. also thanks the financial support from the Estonian Research Council through the personal research grant PUT1476. This work was supported by the Estonian Centre of Excellence in IT (EXCITE), funded by the European Regional Development Fund. C.B. was funded by the Royal Society and Wellcome Trust. We gratefully acknowledge the support of NVIDIA Corporation with the donation of one GeForce GTX TITAN X GPU used for this research. The funders had no role in study design, data collection and analysis, decision to publish, or preparation of the manuscript.

